# Iterative immunostaining combined with expansion microscopy and image processing reveals nanoscopic network organization of nuclear lamina

**DOI:** 10.1101/2022.09.27.509734

**Authors:** Elina Mäntylä, Toni Montonen, Lucio Azzari, Salla Mattola, Markus Hannula, Maija Vihinen-Ranta, Jari Hyttinen, Minnamari Vippola, Alessandro Foi, Soile Nymark, Teemu O. Ihalainen

## Abstract

Investigation of nuclear lamina architecture relies on super-resolved microscopy. However, epitope accessibility, labeling density, and detection precision of individual molecules pose challenges within the molecularly crowded nucleus. We developed iterative indirect immunofluorescence (IT–IF) staining approach combined with expansion microscopy (ExM) and structured illumination microscopy to improve super-resolution microscopy of subnuclear nanostructures like lamins. We prove that ExM is applicable in analyzing highly compacted nuclear multiprotein complexes such as viral capsids and provide technical improvements to ExM method including 3D-printed gel casting equipment. We show that in comparison to conventional immunostaining, IT-IF results in a higher signal-to-background –ratio and a mean fluorescence intensity by improving the labeling density. Moreover, we present a signal processing pipeline for noise estimation, denoising, and deblurring to aid in quantitative image analyses and provide this platform for the microscopy imaging community. Finally, we show the potential of signal-resolved IT–IF in quantitative super-resolution ExM imaging of nuclear lamina and reveal nanoscopic details of the lamin network organization - a prerequisite for studying intranuclear structural co-regulation of cell function and fate. (Words: 175)

## Introduction

Nuclear envelope (NE) is underlined by a tight proteinaceous network called nuclear lamina (NL). The NL is formed of type V intermediate filaments called A and B-type lamins and associated proteins (Schirmer *et al.*, 2003). The A- and B-type lamins form separate layers with the latter situated closer to the NE (Figueiras, Oscar F Silvestre, *et al.*, 2019). Together, lamins form approximately a 14 nm-thick layer less than one-tenth of the resolution of the diffraction-limited fluorescence microscopy (Pawley, 2006). The NL is necessary for the nuclear mechanical stability (Dahl *et al.*, 2004) and responds to mechanical cues (Ankam *et al.*, 2018). It has pivotal roles in regulating genome organization and gene expression (Brueckner *et al.*, 2020) by tethering chromatin (Robson *et al.*, 2016). The NL –chromatin interaction is highly dynamic (Dixon *et al.*, 2015), and changes, e.g., during differentiation (Cavalli and Misteli, 2013; Fortin and Hansen, 2015). To study the role of lamins in the co-regulation of the genome, a detailed understanding of the NL organization is required. Hence, super-resolution imaging of the NL has gained significant interest.

Expansion microscopy (ExM) is a tempting super-resolution microscopy technique enabling the detection of complex nanostructures by using fluorescence-based imaging modalities, e.g., selective plane illumination microscopy or laser scanning confocal microscopy (LSCM). ExM is based on isotropic enlargement of the sample via 4X – 10X expandable hydrogel. However, this expansion dilutes the fluorophores in the sample reducing the resulting image brightness. The dilution scales to the third power of the (linear) expansion factor and thus even 4X expansion can lead to a 64X reduction of the fluorescent signal. To compensate for the dilution, high-quality imaging with ExM, as with other super-resolution methods, requires high-density labeling of the targets (Dankovich and Rizzoli, 2021). However, conventional indirect immunostaining can be insufficient and include non-specific binding, which causes off-target staining of the background and consequent reduction in the image signal-to-background ratio (SBR) and contrast (Lau *et al.*, 2012; Whelan and Bell, 2015). Furthermore, the accessibility of the target protein epitopes can drastically affect the quality of the data (Schnell *et al.*, 2012). The low signal quality in ExM can be improved by post-processing the data by denoising (Chen *et al.*, 2021), deconvolution (Ikoma *et al.*, 2018), or by a combination of both.

We present an improved ExM method utilizing iterative immunofluorescence (IT-IF) staining and image processing further facilitating detection and quantitative analysis in super-resolution microscopy of the nucleus. We provide free image processing tools specifically designed to perform noise reduction on LSCM data. We prove that IT-IF leads to increased signal intensity without compromising the SBR, advancing super-resolution imaging of highly compact intranuclear structures. Finally, we exploit these methods to reveal nanoscopic structural details of NL network organization.

## Results and discussion

### ExM enables high-resolution imaging of highly compacted nuclear multiprotein complexes but compromises fluorescence intensity and signal-to-background-ratio

ExM allows super-resolution imaging of biological specimens using standard fluorescence microscopy systems. It is based on the isotropic physical expansion of the sample, leading to a corresponding increase in the spatial resolution (Tillberg *et al.*, 2016). Here, fixed and immunostained samples are treated with a cross-linker, embedded into polyacrylamide (PAA) hydrogel, homogenized, and osmotically expanded in water (Figure 1A). We investigated if ExM results in isotropic expansion of rigid and compact nano-scale structures within the nucleus. For this, we used HSV-1-infected fibroblasts (12 h post-infection, multiplicity of infection 5) (Figure 1B). HSV-1 naturally targets nuclei for replication producing 125 nm-wide self-assembling progeny capsids in the nucleoplasm (Newcomb *et al.*, 1996; Ojala *et al.*, 2000; Baines, 2011). After immunostaining with a viral capsid protein 5 (VP5) antibody (Ab), the samples were prepared for ExM (n=2, expansion factor ∼4), imaged with LSCM, and deconvolved. From the images, 12 intranuclear anti-VP5 labeled ring-like viral capsids were extracted, aligned, averaged, and their mean diameter was measured as the peak-to-peak distance from the normalized intensity profile. The mean capsid size was 125.4 nm correlating well with that of the native virus (Figure 1B). This is in good agreement with previous studies, where super-resolution STORM-imaging of HSV-1 resulted in approximately 133 nm diameter for the tegument layer outside the capsid (Laine *et al.*, 2015). Thus, the assay showed successful isotropic expansion of the intranuclear viral capsids, indicating ExM suitability for super-resolution imaging of compact intranuclear objects (Gao *et al.*, 2021).

**Figure 1.**
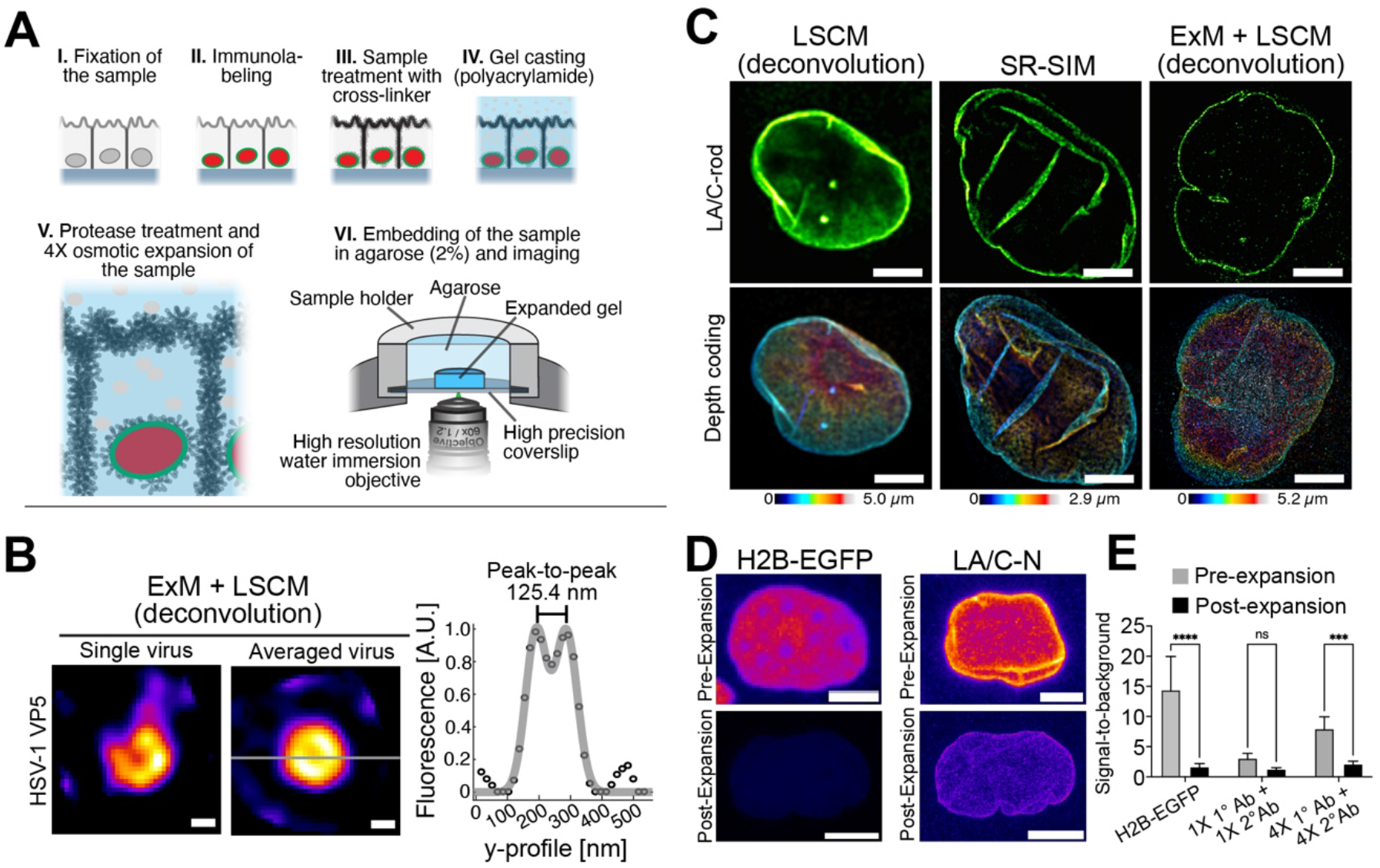
Graphical presentation of pre-expansion microscopy (pre-ExM) sample preparation method and influence of ExM on detection of intranuclear protein complexes and signal fluorescence intensity. (A) Schematics of pre-ExM sample preparation method and imaging setup showing critical steps of sample fixation (i), immunostaining (ii), Acroloyl-X crosslinking (iv), protease treatment followed by the osmotic ∼4X expansion (v), and finally, mounting of the gel sample into an AIREKA Cell for inverted laser scanning confocal microscopy (LSCM) using a high-resolution water immersion objective (vi). (B) Representative deconvoluted post-ExM confocal microscopy images of intranuclear single (left) and an averaged (n=6; right) viral protein 5 (VP5) Ab-stained HSV-1 capsids and the mean fitted capsid diameter (125.4 nm) measured as a peak-to-peak distance from the fluorescence histogram proving the symmetric expansion of even a complex nuclear protein complex. Scale bars, 40 nm. (C) Deconvoluted LSCM (upper right), super-resolution structured illumination microscopy (SR-SIM) (upper middle), and deconvoluted ExM+LSCM images of an epithelial cell nucleus stained with a rod-domain targeting lamin A/C Ab (Ab) and depth-coded pseudo-color fluorescence intensity maps (lower panels) showing the effect on the detection resolution and proving the power of ExM in imaging of nuclear substructures. Scale bars, 5 µm. (D) Representative pseudo-colored pre-ExM (upper panel) and post-ExM (lower panel) LSCM images of endogenously expressed H2B-EGFP and LA/C-N-stained (1X primary (1°) and secondary (2°) Abs) epithelial cell nuclei showing the decrease in fluorescence intensity after the ExM method. Scale bars, 5µm. (E) Quantification of the signal-to-background ratio (SBR) before and after expansion for H2B-EGFP, and LA/C-N staining with conventional (1X) or enhanced (4X) concentrations of both 1° and 2° Abs. Columns represent the measured values as a mean ± standard deviation (ns p>0.05, *** p<0.001, **** p<0.0001, One-way ANOVA multiple comparisons).

We then compared the performances of LSCM, SR-SIM, and ExM+LSCM in the imaging of NL (Figure 1C). Previous cryo-electron tomography and fluorescence lifetime – based studies on the molecular architecture of the NL (∼14 nm) have shown that A-type lamins A and C (LA/C) form a separated network of ∼10 nm in thicknesses (Turgay *et al.*, 2017; Figueiras, Oscar F Silvestre, *et al.*, 2019). Thus, the NL falls beyond the diffraction-limited resolution of a confocal microscope and presents a good intranuclear target for comparison of different imaging modalities. Here, cells were stained with a mouse monoclonal Ab (mMAb) targeting the lamin A/C rod-domain (LA/C-rod) and mounted for high-resolution LSCM and SR-SIM (n=2). Consequently, LA/C-rod –stained ExM samples were prepared from replicate samples by crosslinking and embedding into the PAA gel using novel in-house 3D-printed spacers designed to ensure constant sample gel thickness and diameter (Supplementary Figure 1). After homogenization, the samples were expanded in water (expansion factor ∼4.4, n=2). Following imaging with LSCM, SR-SIM, and ExM+LSCM, the width of the LA/C rim was determined from the middle cross-section of the nucleus by measuring the full width at half maximum (FWHM) (Figure 1C). The LSCM yielded the thickest lamina with a mean thickness of 273.0 nm ± 0.04 nm. The SR-SIM improved the resolution, and the lamina thickness was 182.0 nm ± 0.4 nm. The ExM provided the highest resolution, with a lamina thickness of 110.0 nm ± 0.4 nm. LSCM and SR-SIM showed surprisingly mediocre performance and the resulting thicknesses were much larger than the theoretical diffraction limit. The results indicate that the ExM yielded considerably improved resolution (smaller FWHM of the NL) in comparison to LSCM or SR-SIM when imaging intranuclear structures.

Ab concentrations used in ExM are typically higher than those used in conventional immunostaining protocols because in ExM the fluorescence signal is weakened by a factor of expansion in the power of three (Truckenbrodt *et al.*, 2019). To demonstrate this effect, we next prepared pre- and post-expansion samples from cells stably expressing histone H2B-EGFP. In addition, to visualize the effect of expansion on the Ab-derived intensity, replicate samples were stained with an Ab recognizing LA/C N-terminus (LA/C-N) using standard, i.e., 1X primary (1°) and 1X secondary (2°) Ab concentrations. The fluorescence intensity and the mean SBR were determined with identical LSCM imaging settings before and after the expansion (expansion factor 4.4; n=2). Of note, the SBR here described the ratio of background-corrected nuclear intensity to the intensity resulting from off-target and background staining measured from the cytoplasm. Also, to ensure that the detection conditions did not interfere with the analysis of weak fluorescence intensities, the effects of detector voltage and laser power on the linearity of the detection were determined (Supplementary Figure 2, A and B).

The stably expressed H2B-EGFP intensity was found to be 1620 ± 140 artificial units (a.u.) (mean ± standard deviation, SD) before the expansion but was significantly reduced by 91 % to 153 ± 41 a.u. in the expansion (n=2). Consequently, the SBR before the expansion was 14.7 but decreased by 88 % to 1.7 after the expansion (Figure 1, D and E). For LA/C-N (n=2), the standard labeling-acquired intensity was 257 ± 53 a.u. before the expansion reducing to 183 ± 81 a.u. after the expansion leading to a ∼30 % decrease in the intensity. Consequently, the LA/C-N expansion led to a 58 % decrease in the mean SBR (from SBR of 3.1 to 1.3) (Figure 1E).

Together, the results indicate that ExM enables imaging of highly compacted nuclear protein assemblies. Also, the results describe the reduction in intensity and SBR in ExM imaging demonstrating that the ExM decreases signal intensity due to the dilution of the fluorophores within the sample (Truckenbrodt *et al.*, 2018; Wassie, Zhao and Boyden, 2019; Gaudreau-Lapierre *et al.*, 2021). The caveat here is that proper imaging of samples like these requires longer acquisition times, higher laser power, and higher detector gains leading to increased noise and photobleaching. As our analysis of the pre-and post-ExM gel-embedded samples critically showed, the labeling method requires further development to counteract the resulting decrease in sample fluorescent intensity and signal quality.

### Altering the concentration of primary and secondary Abs influences fluorescence signal intensity and signal-to-background ratio

To understand how the intensity and signal quality could be improved in ExM imaging, we investigated the effect of high Ab concentrations on the intensity and the SBR before and after the ExM. For this, samples were prepared with 4X higher LA/C-N 1° Ab (4X 1°) and 4X higher 2° Ab (4X 2° Ab) concentrations (n=2). The analysis showed that before the expansion, the intensities of samples containing this 4X 1°/4X 2° Ab concentrations were almost 4X higher (950 ± 150 a.u.) in comparison to that acquired with the standard 1X labeling concentrations (257 ± 53 a.u.). Following the expansion, the intensity was decreased by 54 % to 435 ± 72 a.u. Also, the expansion led to a 71 % decrease in the SBR from 6.6 to 1.9 (Figure 1E). However, our results show that increasing the Ab concentration solely does not rescue the image quality. Next, we sought to separately analyze the effect of increased 1^°^ Ab and 2^°^ Ab concentrations on LA/C-N Ab labeling in different microscopy approaches. For this, we first used either 4X 1° Ab or 2° Ab concentration with respect to the standard 1X concentration of the other. The 4X 1^°^ Ab and 4X 2^°^ Ab samples were then imaged with ExM-LSCM (n=2) (Figure 2, A-D), SIM (Figure 2, E and F), and LSCM (Figure 2, G-J) (see also Supplementary Figure 3 for rabbit monoclonal Ab LA/C-rod and rabbit polyclonal Ab H3). As polyclonal antibodies have multiple binding sites with potentially different binding affinities, in the following experiments monoclonal LA/C-N Ab was used as it has a single known binding site specific for an epitope between amino acids 2-29 at the N-terminus of lamin A/C.

**Figure 2.**
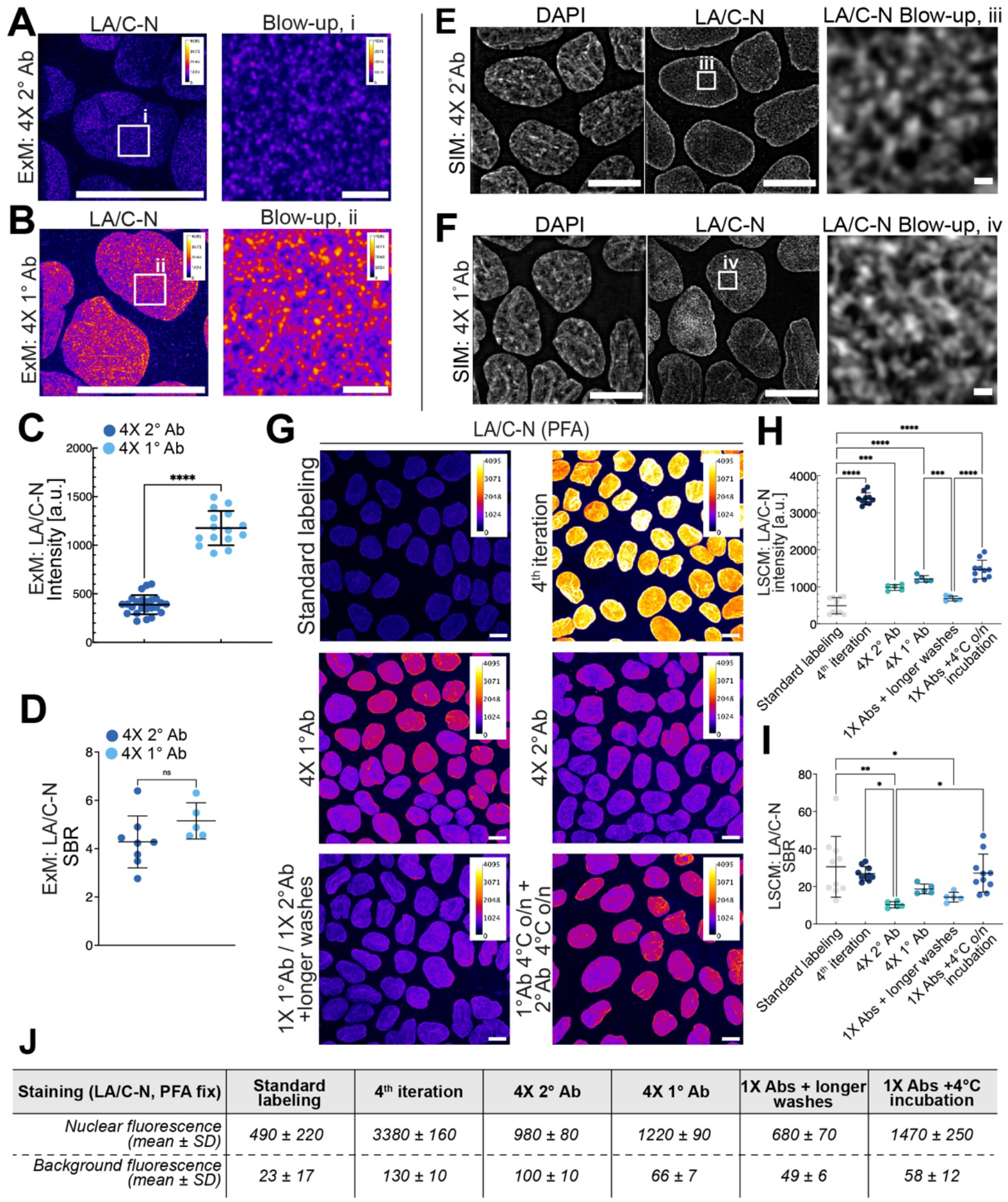
Effects of enhanced primary and secondary Ab concentration on target detection. (A) Representative expansion microscopy (ExM) + LSCM pseudo-colored maximum intensity projection image of 4-times increased secondary Ab concentration (4X 2°Ab) and (B) 4-times increased primary Ab concentration (4X 1°Ab) (in left panels: scale bars, 20 µm; in right panels: scale bars, 1 µm) and their respective blow-ups (i and ii, right panels). (C) Quantifications of LA/C-N intensity and (D) SBR in ExM images of samples treated with 4X 1°Ab and 4X 2°Ab. The data represents mean ± standard deviation (** p<0.01, **** p<0.0001, Student’s T-test) (E) Structured illumination microscopy (SIM) maximum intensity projection images of DAPI and LA/C-N and their respective blow-ups (iii and iv) using 4-times increased secondary Ab concentration (4X 2°Ab) and (F) 4-times increased primary Ab concentration (4X 1°Ab). Scale bars, 20 µm; blow-up scale bars, 1 µm. (G) Representative pseudo-colored LSCM images of LA/C-N-stained epithelial nuclei using either 1X primary (1X 1°Ab) with 1X secondary (1X 2°Ab) concentration i.e., standard labeling (upper left panel), four times iterated (4^th^ iteration) sample (upper right panel), 4-times increased primary Ab concentration (4X 1°Ab, middle left panel), 4-times increased secondary Ab concentration (4X 2°Ab, middle right panel), 1X 1°Ab with 1X 2°Ab treatments with longer washes (all PBS and ddH_2_O washes prolonged to 1h, lower left panel), and overnight (o/n) incubations of both 1X 1°Ab and 4X 2°Abs at 4°C (lower right panel). Scale bars, 10 µm. (H) Quantification of the mean LA/C-N fluorescence intensity and (I) signal-to-background ratio (SBR) in applied combinations of Ab concentrations and treatments. The data represents mean ± standard deviation (*p<0.05, **p<0.01, ***p<0.001, ****p<0.0001, One-way ANOVA, multiple comparisons). (J) Table of the exact values of fluorescence intensities acquired from the nuclei and the background after the different treatments.

The ExM-LSCM imaging revealed that an increased concentration of the 1^°^ Ab was required to obtain a better signal in ExM (Figure 2, A and B). The LA/C-N samples treated with 4X 2^°^ Ab had a low mean intensity (390.0 ± 100.0 a.u., Figure 2, A and C) and low labeling density (Figure 2A). In contrast, the 4X 1^°^ Ab treatment had a significantly increased intensity (1200.0 ± 180.0 a.u.) (Figure 2, B and C) and visually appeared more densely stained with improved detection of filamentous lamin structures (Figure 2 B). Furthermore, 4X 1^°^ Ab treatment slightly but non-significantly improved the mean SBR (5.0 ± 1.0) (Figure 2D) when compared to 4X 2^°^ Ab treatment (4.0 ± 1.0, Figure 2D). Of note, similar ExM results were also obtained with H3 Ab, where 4X 1^°^ Ab treatment resulted in a significantly higher intensity (1400 ± 250 a.u.) and SBR (1.5 ± 0.1) in comparison to those acquired with 4X 2^°^ Ab treatment (99 ± 59 a.u. and 1.3 ± 0.2, respectively) (Supplementary Figure 3, A and B).

The SIM imaging of the LA/C-N samples showed a better-resolved lamin network in the 4X 1^°^ Ab-treated sample in comparison to the 1X 1^°^ Ab treated one (Figure 2, E and F). However, the treatments did not influence the exposure time during SIM imaging. In line with this, SIM imaging of H3 Ab also showed improved detection with the 4X 1^°^ Ab in comparison to the 4X 2^°^ Ab treatment (Supplementary Figure 3, C and D) while having similar SIM exposure times.

These experiments suggested that in ExM and SIM the resulting intensity and SBR depended on the used Ab concentrations potentially affecting the Ab abundance in the sample. Thus, we sought to study in more detail how other labeling parameters influence the resulting intensity and SBR. We used LA/C-N Ab in LSCM and compared the intensities and SBRs acquired with either standard immunostaining (1X 1° and 2° Abs), concentration enhancement (4X 1^°^ Ab or 4X 2^°^ Ab treatments), prolonged washes, and over-night incubations at +4°C (n=2) (Figure 2, G-J). The analysis of samples stained with the standard immunostaining had a mean nuclear fluorescence intensity of 490 ± 220 a.u. with a mean SBR of 30 ± 20. In comparison, using 4X 1^°^ Ab or 4X 2^°^ Ab improved the fluorescence intensity (1220 ± 90 a.u.) (Figure 2, G-H and J). However, both treatments resulted in notably lower mean SBRs (19 ± 3; ns, and 10 ± 2) in comparison to the traditional staining (Figure 2I). These analyses showed that with the mouse monoclonal LA/C-N, increasing the concentration of 1^°^ Ab in relation to the 2^°^ Ab seemed to be the more crucial factor in improving the sample intensity in LSCM. However, rabbit monoclonal LA/C-rod and rabbit polyclonal H3 Ab produced contradictious results and indicated that there the 4X 2^°^ Ab improved the fluorescence intensity (Supplementary Figure 3, E-H). With 4X 2^°^ Ab the nuclear LA/C-rod and H3 intensities were 410 ± 9 a.u. and 120 ± 20, whereas those in the 4X 1^°^ Ab treatment were 130 ± 9 and 60 ± 20 a.u., respectively. The 4X 2^°^ Ab treatment significantly improved the LA/C-rod SBR (6.0 ± 0.4) in comparison to the 4X a^°^ Ab (3.0 ± 1.0). However, this treatment did not affect the H3 SBR. While the SBR in 4X 1^°^ Ab treatment for H3 Ab was 1.0 ± 0.2, the SBR in the 4X 2^°^ Ab-treated sample was equal (1.0 ± 0.1) with no statistically significant difference. We also found with LA/C-N Ab that using prolonged washes did not significantly affect the intensity (680 ± 70 a.u.), but +4°C incubation with both 1° and 2° Abs significantly improved it (1470 ± 250 a.u.) in comparison to the standard IF staining. However, none of these treatments improved the SBR (Figure 2, G-H and J). However, it should be noted that the staining outcome from altering the Ab concentration ratio is presumably Ab type and affinity-dependent, and we highlight the importance of optimizing the labeling conditions. In the end, the optimal Ab concentration is a balance between the affinity of the Ab for its targeted epitope and its non-specific binding.

Finally, we hypothesized that the antibody epitope accessibility might influence the labeling outcome and affect the intensity. Thus, it was investigated if repetitive labeling (iterative standard immunolabeling) would increase the image quality. Interestingly, our experiments showed that especially the sample intensity can be substantially improved by labeling the sample several times. The four-times iterated (4^th^ iteration) sample had a significantly increased and highest mean fluorescence intensity (3380 ± 160 a.u.) when compared to 4X 1° or 4X 2° Ab samples (Figure 2, G and H). In concert, the SBR in 4^th^ iteration was found to be 27 ± 4 (Figure 2I), indicating that despite the marked increase in intensity, this treatment did not compromise the SBR in comparison to the traditional immunostaining.

### Iterative indirect immunostaining improves fluorescence intensity and signal-to background-ratio enhancing imaging properties in LSCM, SIM and especially in ExM

To study further the possibilities of iterative staining, we developed an iterative indirect immunostaining (IT-IF) protocol. Here, subsequent immunostaining cycles were performed four times, omitting the blocking step in the last three. Each round involved four steps: i) permeabilization followed by ii) incubation with a 1^°^ Ab with uniquely determined optimal concentrations (referred as 1X), iii) washes, and finally, iv) 2^°^ Ab incubation by using optimized concentrations followed by LSCM imaging.

For the iteration experiments, the nuclear lamin A/C was stained with either monoclonal Abs targeting its N- (LA/C-N, PFA and MetOH fixed), C-terminus (LA/C-C, PFA fixed) or rod-domain (LA/C-rod, PFA fixed) Ab, or a rabbit polyclonal Ab (pLA/C, MetOH fixed) (Figure 3). Histone H3 and H3K9me3 Abs and fluorophore-conjugated phalloidin against F-actin were used as controls (Supplementary Figure 4).

**Figure 3.**
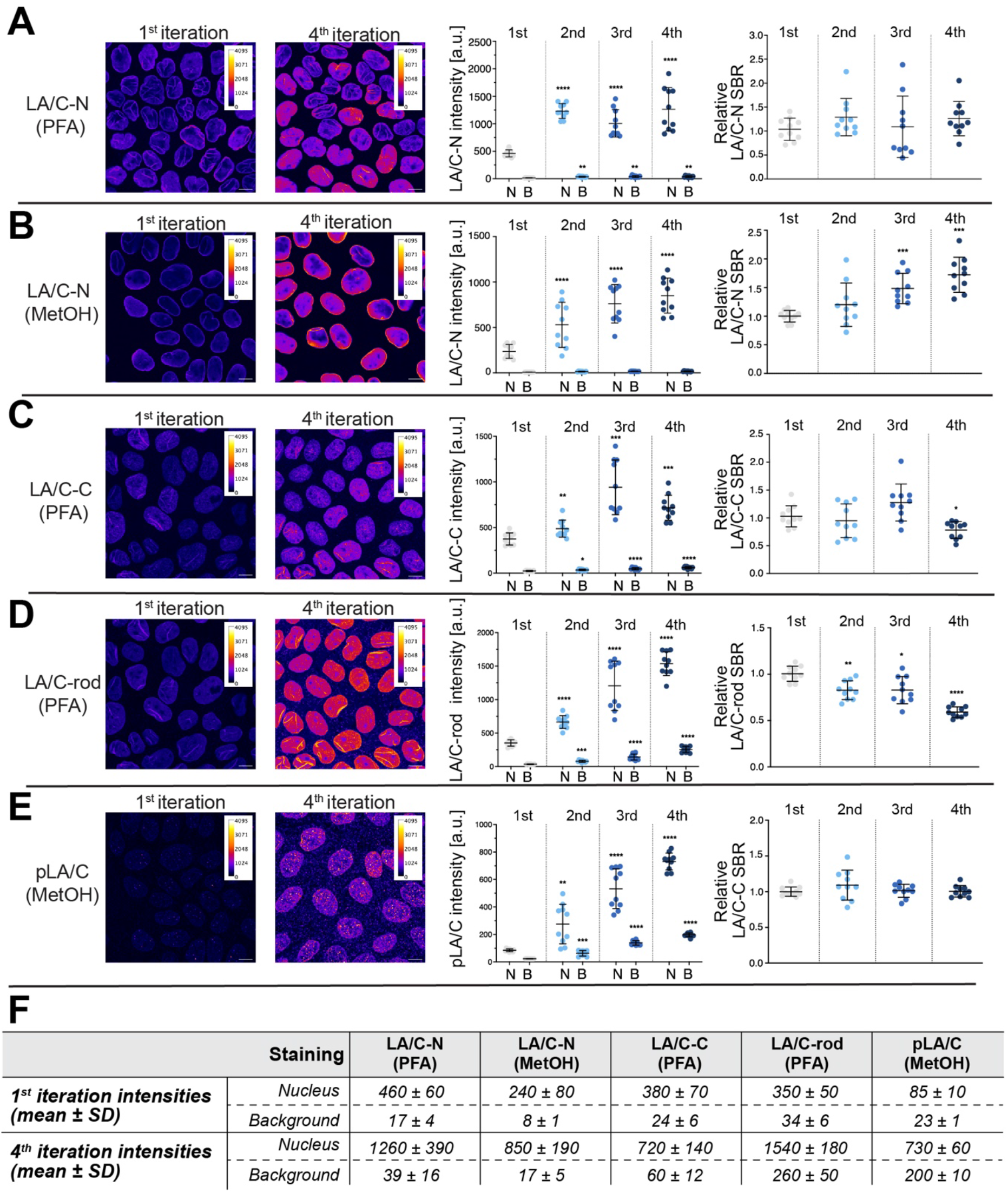
Effect of iterative immunostaining on fluorescence signal intensity and detection quality of nuclear lamins. Representative pseudo-colored maximum intensity projections of LSCM data acquired after one (far left panels) or four (left panels) iterations i.e., consequent immunostaining cycles, and their respective quantifications of background-corrected mean fluorescence intensity (middle right panels), and background-corrected signal-to-background ratio (SBR, normalized to iteration 1, far right panels) of (A) LA/C-N on PFA-, (B) LA/C-N on MetOH-, (C) LA/C-C on PFA, (D) LA/C-rod on PFA-, and (E) pLA/C on MetOH-fixed cells. Scale bars, 10 µm. F) Table containing the exact values from the above measurements of nuclear and background fluorescence intensities. All values are presented as a mean ± standard deviation (ns p>0.05, *p<0.05, **p<0.01, ***p<0.001, ****p<0.0001, One-way ANOVA, multiple comparisons, data compared to the 1^st^ iteration).

The analyses of normalized intensity values (see Materials and Methods) showed that the total fluorescence intensity was increased due to the iterative staining with all tested Abs but not with the phalloidin (Figure 3, Supplementary Figure 4). For the lamin-stained samples, the iterations increased the background-corrected mean fluorescence intensity of LA/C-N by ∼2.7 fold in PFA-fixed (Figure 3A) and by ∼3.6 fold in MetOH-fixed samples (Figure 3B). Similarly, the intensities were increased in PFA-fixed LA/C-C (by ∼1.9 fold) and LA/C-rod (∼4.4 fold) stained, and MetOH-fixed pLA/C (by ∼8.6 fold) stained samples (Figure 3, C-F). Consequently, the background intensity (I_B_) was increased by ∼2.4 fold for PFA-fixed LA/C-N, by ∼2.2 fold for MetOH-fixed LA/C-N, by ∼2.5 fold for PFA-fixed LA/C-C, by ∼7.4 fold for PFA-fixed LA/C-rod, and lastly, by ∼8.6 fold for MetOH-fixed pLA/C (Figure 3, A-F). However, even after four iterations, the background staining interfered neither with the detection nor the visual interpretation of the results with the studied Abs. This was apparent, especially from the quantitated SBRs. Specifically, the mean background-corrected SBRs after the first iteration were ∼28.80 for LA/C-N (PFA), ∼31.42 for LA/C-N (MetOH), ∼16.26 for LA/C-C (PFA), ∼10.37 for LA/C-rod (PFA), and ∼3.73 for pLA/C (MetOH) (Figure 3, A-E, right panels). In addition, the H3 (PFA) mean relative intensity increased between 1^st^ and 4^th^ iteration by ∼8.9 fold, and for MetOH-fixed H3K9me3 the mean relative intensity increased from 1^st^ to 4^th^ iteration by ∼6.2 fold (n=2) (Supplementary Figure 4). To analyze and compare the SBRs in the iterations, the SBRs for the following iterations were normalized to that from the 1^st^ iteration) (Figure 3, A-E, right panels). After the four iterations, the relative SBR in LA/C-N (PFA) staining remained nearly unaltered (from 1 to 1.26, statistically non-significant, ns, p>0.05). In contrast, in MetOH fixed LA/C-N samples, the relative SBR was increased by each iteration and was significantly improved after the fourth iteration (from 1 to 1.7). For the LA/C-C (PFA), the relative SBR also remained nearly similar during the first three iterations but slightly decreased after the fourth (from 1 to 0.78). In LA/C-rod (PFA) treated samples, the relative SBR was significantly lowered after the four iterations (from 1 to 0.592). Lastly, the SBR was ∼1.0 in all four pLA/C (MetOH) iterations (*p>0.05). Lastly, the iterations improved slightly but non-significantly the relative SBR of H3 (by ∼1.2 fold, ns), but lowered the SBR of MetOH-fixed H3K9me3 by ∼0,7 fold (n=2) (Supplementary Figure 4, A-C). These analyses show that the iterations increase the intensity of especially PFA-fixed samples and maintain the SBR depending on the Ab specificity. Thus, the absolute difference between the signal and background increased considerably.

It was then sought to determine how the iterations affected the super-resolution imaging of the samples. To this end, SIM and ExM-LSCM imaging were done after the first and fourth iterations. The experiment showed that the iterations enabled SIM imaging with shorter exposure times as the sample intensity was increased. Also, the structural details, including the NL at the nuclear rim, appeared more uniform (Figure 4, A and B). Similarly, the ExM-LSCM imaging indicated that the iterations seemed to increase the sample intensity, and notably, the lamin meshwork appeared more evident and visible (Figure 4, C and D). Specifically, the mean intensity increased significantly by 4-fold following the four iterations from 240 ± 24.0 a.u. of the 1^st^ iteration to 980 ± 180.0 a.u. in the 4^th^ iteration (Figure 4E). Concomitantly, the SBR was also increased by 4-fold from 1.7 ± 0.17 of the 1^st^ iteration to 6.4 ± 1.2 (Figure 4F). Of note, SIM-imaging indicated that the iterations improved imaging of structural details with H3 but did not significantly affect detection with phalloidin (Supplementary Figure 4, D - G). Together, these results indicated that iteration-enhanced labeling density within the sample leads to enhanced intensity, contrast, and structural details.

**Figure 4.**
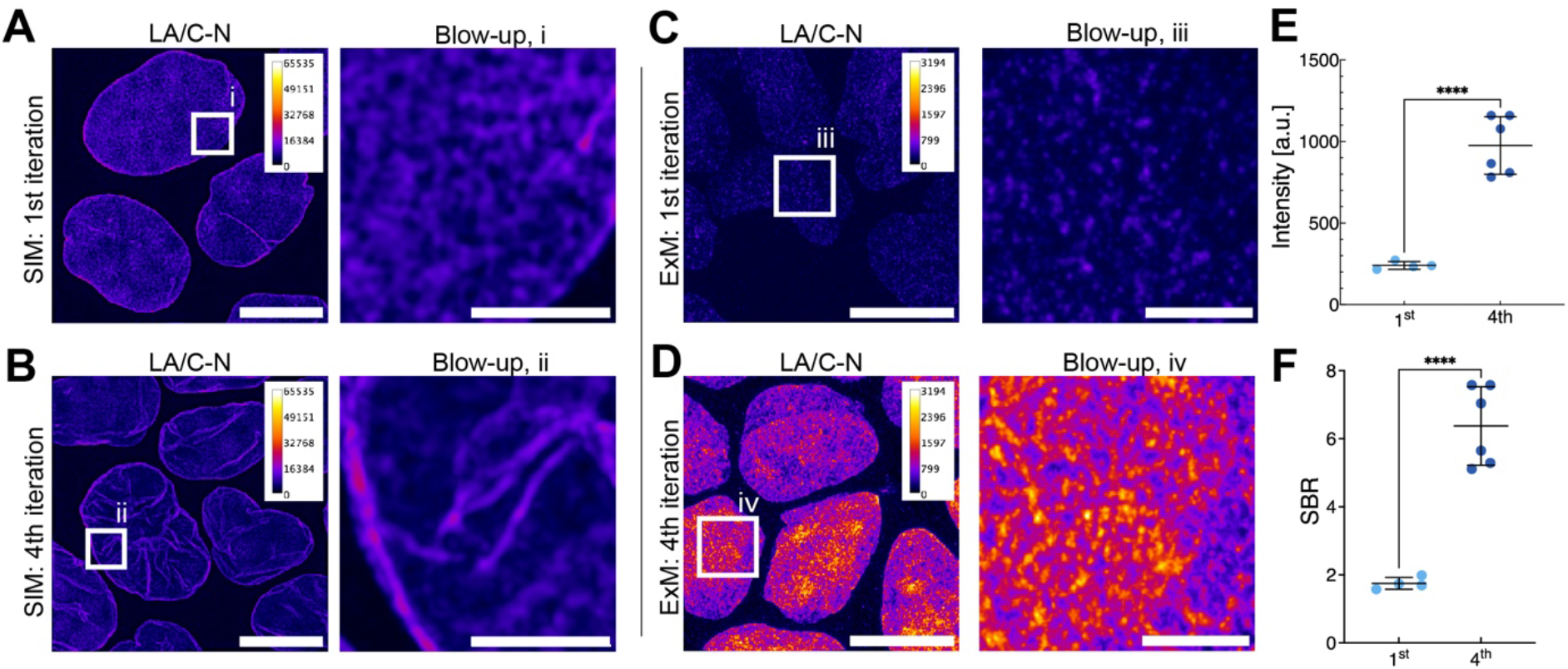
Super-resolution SIM imaging of IT-IF samples. Representative pseudo-colored maximum intensity projections of SIM data and their blow-ups from indicated areas (white box) of (A) first (1^st^) and (B) four times iterated (4^th^) LA/C-N staining. Representative pseudo-colored maximum intensity projections of ExM-LSCM data and their blow-ups from indicated areas (white box) of (C) first (1^st^) and (D) four times iterated (4^th^) LA/C-N staining showing increased intensity and staining density following the 4^th^ iteration. Scale bars, 10 µm; blow-up scale bars, 2 µm; expansion factor 4.4. (E) Quantification of mean fluorescent intensity and (F) signal-to-background ratio (SBR) in 1^st^ and 4^th^ iterations. The data represents mean ± standard deviation (**** p<0.0001, Student’s T-test).

Together, these results show that the IT-IF improves the sample imaging properties in LSCM, SIM and especially in ExM. Specifically, the signal intensity was increased without compromising the SBR. However, it should be noted that while using either low concentrations or low-affinity Abs might reduce background in the conventional microscopy, in ExM it causes low signal intensity to be lost in the noise.

### IT-IF with modeling-based denoising improves quantitative lamin network detection

We sought to determine how the IT-IF improves the analysis of NL structural organization. For this, multicolor detection of the four iterations with LA/C-N was done by using Alexa 647-linked 2°Ab in first and second iterations, and Alexa 568- and Alexa 488-linked 2°Abs in third and fourth iterations, respectively (Figure 5, A and B). This multicolor staining enabled the analysis of labeling patterns after each iteration (Figure 5C) and analysis of their spatial correlations (Figure 5D).

**Figure 5.**
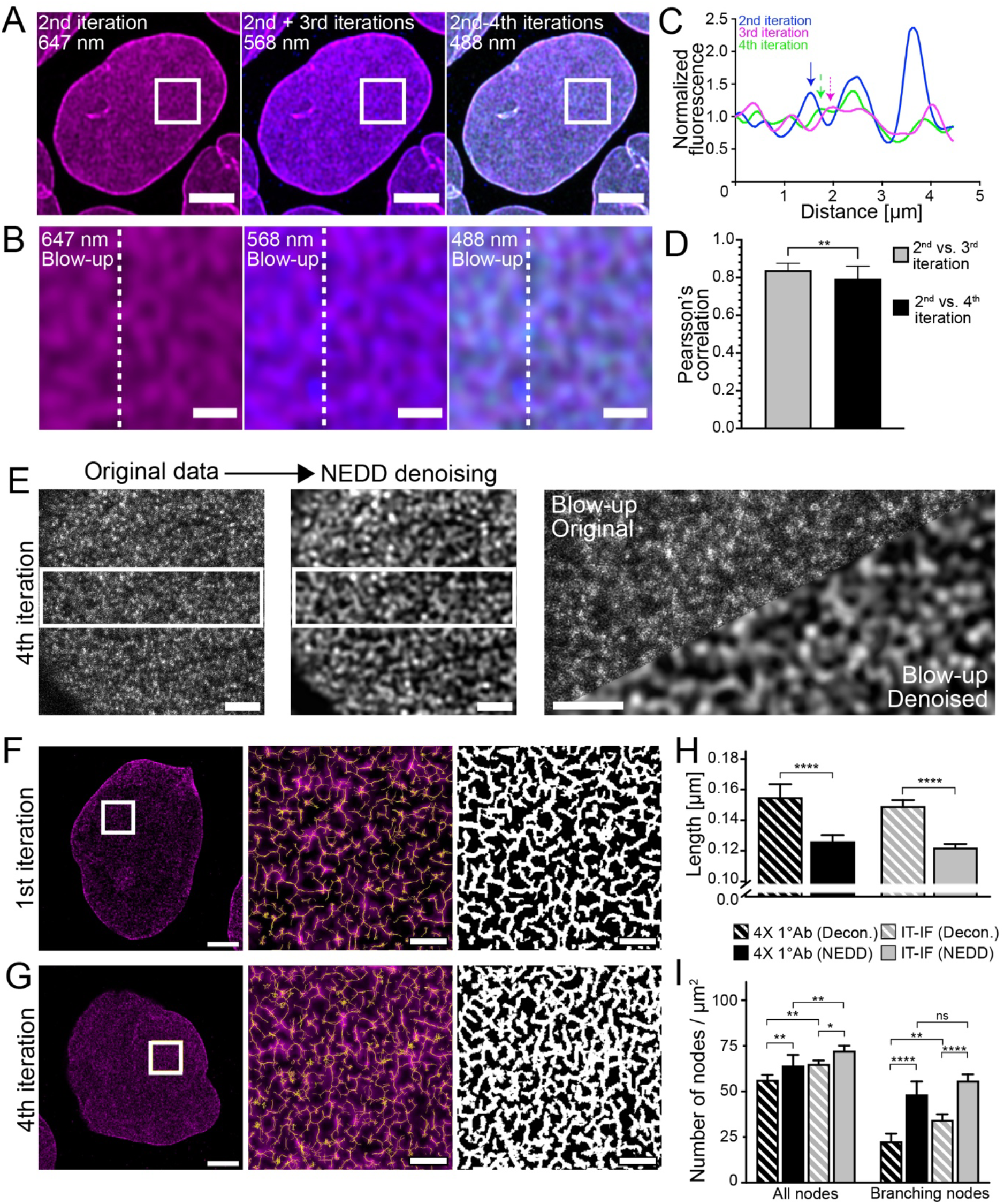
NL network architecture visualized by IT-IF combined with modeling-based denoising. (A) Representative LSCM single optical section images of LA/C-N iteration showing detection result after the first two iterations (Alexa 647, upper left panel, magenta), after 3^rd^ (Alexa 568, upper middle panel, blue), and 4^th^ iteration (Alexa 488, upper right panel, green). Scale bars, 5 µm. (B) Blow-ups of indicated areas shown in A. Scale bars, 1 µm. (C) Fluorescence intensity line profile along the lines indicated in B, showing the fluorescence intensity distribution of combined 1^st^ and 2^nd^ (magenta), 3^rd^ (blue) and 4th iterations (green). Arrows indicate increased Ab binding as iteration-specific intensity peaks. (D) Pearson Correlation Coefficient between indicated iterations. The data represents mean ± standard deviation (** p<0.01, Student’s T-test) (E) NEDD denoising of IT-IF data along with blow-up images side-by-side comparison of the result. Scale bars, 1 µm. Representative NEDD denoised ExM images of single nuclei showing (F) ExM optimized LA/C-N staining using 4X 1°Ab concentrations or (G) four times iterated LA/C-N (left panels, scale bars, 3 µm), blow-up overlay of indicated areas (white boxes) with segmented network structure (yellow) (middle panels, scale bars, 500 nm) and reconstructed LA/C-N network (right panels, scale bars, 500 nm). (H) Quantification of lamin segment length after 4X 1°Ab or IT-IF labeling following deconvolution or NEDD denoising. The data represents mean ± standard deviation (**** p<0.0001, Ordinary One-way ANOVA, multiple comparisons). (I) Quantification of NL architecture by presenting the number of total and branching nodes. The data represents mean ± standard deviation (ns, p>0.05, * p<0.05, ** p<0.01 **** p<0.0001, Ordinary One-way ANOVA, multiple comparisons).

The multicolor analysis revealed that each iteration improved the detection efficiency of lamin by enhancing the Ab occupancy through binding to new targets not bound during preceding iterations (Figure 5, A and B). This was confirmed by intensity line profile analysis showing iteration-specific intensity peaks, and by Pearsońs correlation coefficient (PCC) analysis where the correlation between the stainings was reasonably strong but weakened significantly during the consecutive iterations (Figure 5, C and D, n=2). The PCC after the two first iterations in comparison to the third was found to be 0.839 ± 0.037, while being significantly lowered to 0.794 ± 0.067 in comparison to that of the fourth iteration (n=3) (Figure 5D). These results suggested that each consecutive iteration produced new and additional Ab binding along the lamins.

Finally, the power of IT-IF in the detection of lamin network organization was studied by comparing 1°Ab concentration-enhanced (4X 1^°^ Ab) and four times iterated LA/C-N – stained IT-IF samples by using ExM (n=2). Again, the measured data featured a low signal quality highlighting the need for solutions to get a faithful estimation of the clean signal for reliable quantification of the network properties. For this, we developed a signal-processing software for Noise Estimation, Denoising, and Deblurring (NEDD) [https://github.com/lucioazzari/NoiseEstimationDenoisingDeblurring] that automatically estimates the noise and performs signal reconstruction intended as denoising and deblurring (see Materials and Methods for detailed description for the pipeline). After processing the images with NEDD, we qualitatively observed that in comparison to the concentration enhanced-samples, the four-time iterated IT-IF samples, displayed a clear improvement in the overall signal quality showing well-defined lamin structures (Figure 5 F and G). Specifically, IT-IF ExM enabled detailed quantitative analysis of the lamin network architecture. The physical properties of the LA/C-N –stained lamins were further quantified by analyzing the lamin segment length and the NEDD method was compared to traditional deconvolution based denoising. The analysis showed similar segment length for both 4X 1°Ab and IT-IF samples after deconvolution (0.16 ± 0.01 µm and 0.150 ± 0.005, respectively). However, NEDD lead to reduced length of the quantified segment length (0.126 ± 0.005 µm and 0.122 ± 0.003, respectively) when compared to deconvolved data (Figure 5H). Previous cryo-electron tomography imaging revealed that individual lamin filaments are approximately 380 nm in length (Turgay et al. 2017). In line with this, super-resolution dSTORM imaging studies have indicated roughly 150 nm lamin segment length in the dense lamina network (Xie *et al.*, 2016), which is highly similar to the segment length quantified here by using IT-IF ExM.

To further analyze the LA/C network properties after IT-IF, we next quantified the total number of nodes and number of terminating nodes per square micrometer of the nuclear lamina. The analysis indicated a significantly higher total number of nodes in the four times iterated IT-IF samples when compared to 4X 1°Ab after deconvolution (65 ± 2 and 56 ± 3 nodes per µm^2^). The total number of nodes was overall higher after NEDD, indicating again better separation between adjacent nodes (IT-IF vs. 4X 1° Ab, 72 ± 3 and 64 ± 6 nodes per µm^2^, respectively) (Figure 5I). Similarly, the number of detected branching nodes followed the same trend, increasing from 4X 1° Ab to IT-IF (deconvolution data, 23 ± 5 and 34 ± 4 nodes per µm^2^, respectively) and from deconvolution to NEDD (IT-IF data, 34 ± 4 and 56 ± 4 nodes per µm^2^, respectively) (Figure 5I).

Together, the results indicate that the IT-IF and NEDD improved the detection and quantification of the NL structural organization in ExM. The lamin A/C organization revealed here by IT-IF ExM is similar to previous studies conducted with SR-SIM (Shimi *et al.*, 2015) or dSTORM and PALM imaging (Xie *et al.*, 2016; Nmezi *et al.*, 2019). In the future it will interesting to investigate the A-type and B-type lamin networks (Figueiras *et al.*, 2019; Nmezi *et al.*, 2019) and their changes in different pathologies by IT-IF ExM imaging. Here especially the usage of 10X expansion microscopy methods (Damstra *et al.*, 2021; Klimas *et al.*, 2023) would allow better separation of the different networks and their layered structures when compared to original 4X ExM. Finally, antibody epitopes in lamin proteins can be differentially accessible (Ihalainen *et al.*, 2015; Wallace *et al.*, 2023). We speculate that in IT-IF the exposure of buried epitopes is the primary factor contributing to the increase in signal intensity and the resulting density of the detected lamina network.

Overall, the findings of this study indicate that IT-IF combined with ExM and signal processing advance the detection of nuclear lamin structure in the super-resolution scale by providing high labeling density, intensity, and uncompromised SBR. Together with our signal processing pipeline, IT-IF revealed a more complete structural organization of nuclear lamins and enabled detailed quantification of the NL network properties. Beyond the NL study presented here, we expect that our IT-IF method and signal processing platform can be useful in a wide range of nanoscopy applications.

## Materials and methods

### Cells and viruses

Madin-Darby canine kidney (MDCK) type IIG cells were maintained in low glucose MEM (#41090093, Thermo Fisher Scientific, Gibco^TM^, Paisley, UK) supplemented with 1 % (vol/vol) penicillin-streptomycin antibiotics (#15140122, Thermo Fisher Scientific) and with 10% fetal bovine serum (#10500064, Thermo Fisher Scientific). Cells were maintained under a standard 37 °C and humidified atmosphere with 5 % CO2, passaged once a week. For the experiments, cells were seeded for seven days on collagen I (#A1064401, Thermo Fisher Scientific) coated cover glasses (18×18 mm for conventional LSCM, 22×22mm for ExM, high performance, D=0.17 mm, Carl Zeiss Microscopy, NY, USA) prior fixation.

For HSV-1 ExM studies, Vero cells (ATCC) grown on glass coverslips in low-glucose DMEM GlutaMAX (#11570586, Gibco, Thermo Fisher Scientific) to 90 % confluency and inoculated with the virus (HSV-1, MOI 5). The infected and noninfected cells were washed with phosphate-buffered saline (PBS) and fixed with 4 % PFA after 17 h post-infection (pi).

### Antibodies

HSV-1 capsids were detected with HSV-1 VP5 MAb (sc-13525, Santa Cruz Biotechnology, Dallas, TX, USA). To detect A-type lamins in this study, mouse monoclonal Abs (mMAbs) against lamin A/C C-terminus residues aa 319-566 (LA/C-C, [131C3], ab18984, Abcam, Cambridge, UK) and lamin A/C N-terminus residues aa 2-29 (LA/C-N, [E-1], sc-376248, Santa Cruz Biotechnology, DA, USA), a rabbit monoclonal Ab (rMAb) against full-length lamin A/C (LA/C-rod, [EP4520-16], ab133256, Abcam), and a rabbit polyclonal Ab (rPAb) against LA/C (exact target sequence declared as proprietary by the manufacturer, ab227176, Abcam) were used. To detect histone H3, a rPAb (exact target sequence declared as proprietary by the manufacturer H3, ab1791, Abcam) was used. rPAb against H3 tail modification at N-terminal aa 1-100 tri-methyl lysine 9 (H3K9me3, ab8898, Abcam) and anti-actin fluorophore-conjugated phalloidin (ATTO-TEC, NY, USA) were applied in Supplementary studies.

### Iterative immunostaining

MDCKII cells were fixed with 4 % paraformaldehyde (cat. No. 15713-S, Electron Microscopy Sciences, Hatfield, PA, USA) for 10 min at room temperature (RT), rinsed twice with 1X PBS, and permeabilized for 10 min in RT using 0.5 % Triton X-100 in PBS supplemented with 0.5 % BSA (Bovine Serum Albumin) (permeabilization buffer A). Iterative immunostaining was initiated by incubating the samples with primary Abs (1°Ab; see Table 1) diluted in PBS containing 3 % bovine serum albumin (BSA) for continuous blocking (1h in RT) using recommended immunofluorescence labeling concentrations (1X condition). Incubation was followed by washing once with permeabilization buffer A, once with PBS, and again with permeabilization buffer A. The incubation-wash cycle was repeated three times to iterate the 1°Ab binding on the sample. However, after the 1^st^ iteration, the subsequent iterations were permeabilized by using 0.2 % Triton X-100 in PBS supplemented with 0.5 % BSA (permeabilization buffer B). After the four iterations, the samples were treated with secondary Alexa-488 and -568 -conjugated goat anti-mouse and anti-rabbit Abs (2° Ab; 1:200 (1X condition) in 3 % BSA/PBS, Thermo Fisher Scientific, Waltham, MA, USA), respectively, for 1h in RT in the dark, followed by one wash in PBS and one in DI water (á 10 min, RT, in dark). Samples were mounted in ProLong Diamond Antifade Mountant with DAPI (Thermo Fisher Scientific) and left to cure at RT in dark o/n before storing at +4 °C before microscopy. For these analyses, n=2-3 replicate experiments containing 5 random image fields with approx. 20 nuclei were used.

For the expansion of iteratively immunostained samples, images from the 1^st^ and 4^th^ iterations were collected from staining with 1X 1°Ab/1X 2°Ab combination iterated once or four times, respectively. After the isotropic expansion, a piece of gel was cut and mounted with 1 % low-melt agarose into an imaging chamber (SC15022, Aireka Cell, Hong Kong, China).

### Expansion microscopy

Sample expansion was performed using the previously described proExM protocol for cultured cells (Tillberg *et al.*, 2016). The samples were first fixed and stained according to the iterative immunostaining protocol described previously. After the immunolabeling, the samples were incubated in an anchoring solution of 1% 6-((acryloyl)amino) hexanoic acid, succinimidyl ester (#A20770, Acroloyl X-SE, Thermo Fisher Scientific) in PBS for over 6 hours in RT. After the anchoring, the samples were washed with PBS for 2 x 15 mins before proceeding to gelation. Before the gelation, glass coverslips (18mm x 18mm) were prepared to work as a base to cast the gel. A piece of parafilm (P7793, Merck) was glued on top of a coverslip with a drop of cyanoacrylate glue (Loctite Super Glue Power Flex Gel Control, Henkel Norden AB, Bromma, Sweden). The parafilm-coated coverslip was then placed on a 6-well plate lid, parafilm side on top, and the parafilm cover paper was removed. Then, an in-house designed 3D-printed spacer was placed on top of the parafilm. The purpose of the spacer is to confine the gel to known dimensions and keep the thickness of the gel as thin as possible. These features will help define the expansion factor and speed up the expansion of the gel.

The gelation solution was prepared on ice by adding the inhibitor (4-Hydroxytempo) and the accelerator (TEMED) to the monomer solution. The initiator (APS) was added to the solution just before pipetting as the solution will gelatinize rapidly after the inclusion of APS. Thus, it is advisable to prepare the solution into aliquots to separate 1,5ml centrifuge tubes (S1615-5500, Starlab International, Hamburg, Germany) for one to two samples at a time. After adding APS, the solution was vortexed and administered in the middle of the opening in the spacer. Then, the sample was placed carefully on top of the gelling solution sample-side towards the gel. A channel in the spacer ensures that excess gelling solution will flow out. A metal nut was placed on the constructed gel mold to provide extra weight. The sample was left to polymerize for 30-45 mins in RT.

After the gel had polymerized, the parafilm-coated coverslip was removed carefully. The gel will not stick to the parafilm, and it is straightforward to remove it. A scalpel can be used to trim excess gel from the edges of the gel button. The gel was placed on a clean 6-well plate (#150239, Thermo-Fisher Scientific) with the sample coverslip still attached. Only one sample was placed per well. It is essential to measure and draw the outlines of the gel button at this time because the gel will start swelling during digestion. It is crucial to know the original size of the gel button when defining the final expansion factor for the sample. However, as we are using custom-made spacers with a well-defined area for the gel button, it will be the same size every time.

The polymerized gel, together with the sample, was immersed in a 10-fold volume of digestion solution with 8 U/mL Proteinase K (P8107S, New England Biolabs, Ipswich, USA) in digestion buffer containing 50 mM Tris pH 8.0, 1 mM EDTA (E5134, Merck), 0.5 % Triton X-100 (T8787, Merck) and 800 mM guanidine HCl (G3272, Merck) in double-distilled water (ddH_2_O). The sample was incubated in the solution o/n at RT in the dark. The gel will detach from the coverslip during incubation or, at the latest, during the first wash with ddH_2_O. The detached gel was moved to a 6-cm dish preserving the vertical orientation of the gel. It is essential to know the side which includes the sample due to the vast size increase of the sample during the next step. The gel was washed with an excess volume of ddH_2_O for 1 hour, 3-5 times. The hydrophilic gel will swell during these washes until the expansion reaches a plateau after the third or fourth wash.

Finally, the expanded gel sample was measured for the expansion factor (final dimensions/ original dimensions). The gel was cut with a custom-made puncher, and the resulting gel disc was mounted in a live imaging chamber (Aireka Cells) with low-melt agarose (A7431, Merck). The chamber was filled with ddH_2_O to prevent sample shrinkage and drying. For these assays n=2-3 independent replicate experiments all containing duplicate gel samples with 10 nuclei.

### Confocal Microscopy

For LSCM, all samples were prepared as two replicates (n=2) and imaged using constant laser powers and detection voltages, enabling comparable quantitative analysis of the fluorescence intensities. At the confocal microscope, Alexa 488 and Alexa 568 were excited with 488 nm and 561 nm solid-state lasers, respectively. Fluorescence was detected with 525/50 nm and 595/50 band-pass filters. For unexpanded samples, Nikon Apo 60x Oil λS DIC N2, numerical aperture (N.A.) 1.40, the working distance (WD) 0.14 mm was used in imaging, and stacks of 1024×1024 pixels were collected with a pixel size of 41-52nm, or 104 nm in the x- and y – directions, and 150 nm in the z-direction. For expansion microscopy sample imaging, Nikon CFI Plan Apo IR SR 60x WI, N.A. 1.27, WD 0.17 mm objective was used. Stacks of 1024×1024 pixels were collected with a pixel size of 23.6 nm (103.8 nm/expansion factor 4.4) in the x- and y-directions and 180 nm in the z-direction. The imaging was done using constant laser powers and detection voltages, enabling a comparative analysis of the fluorescent intensity in each channel. Correct laser alignment was ensured by using sub-resolution fluorescent beads (PS-Speck, Thermo Fisher Scientific). Image analysis was done with ImageJ FIJI distribution (Schindelin *et al.*, 2012). For these the n=2 independent replicate experiments per Ab containing 10 images with approx. 20 nuclei per image.

### Data normalization and analysis of nuclear lamin intensity and colocalization

After imaging, the mean fluorescence intensity of the NL (measured from the total segmented nuclear area) and the background staining intensity (image area omitting the segmented nuclei) were measured. Additionally, detector noise background correction was imaged from similarly mounted samples containing fluorescent beads (from the area omitting the beads). The detector background was then subtracted from the measured values (background correction), and the background-corrected intensity value was normalized to the starting mean intensity value obtained from iteration 1. For Pearsońs correlation coefficient analysis of colocalization in the multicolor experiment, the correct channel alignment was ensured by using sub-resolution fluorescent beads (PS-Speck, Thermo Fisher Scientific). The PCC analysis was done in ImageJ Fiji with the JACoP plugin (Bolte and Cordelières, 2006). n=2 independent experiments.

### Signal processing pipeline for noise estimation, denoising, and deblurring

In our model for confocal microscopy data, we assume the noise to be additive with signal-dependent variance and spatially correlated (Azzariet al., 2018). The signal-dependent variance models the conversion of light into electric charge and the dark thermal noise. Specifically, the noise variance is an affine function of the signal-expectation (Azzari and Foi, 2014). The noise correlation models the optics of the system and the light’s diffraction pattern of light, commonly referred to as point spread function (PSF), after passing through the lenses of the microscope (Cole et al., 2011). We model correlated noise as the convolution between a kernel and a white Gaussian random field. Alternatively, noise correlation can be modeled with the noise power spectral density (PSD), that is the distribution of the noise energy (or variance) in the Fourier domain. Kernel and PSD are related to each other; in particular, the kernel can be calculated as the inverse Fourier transform of the square root of the noise of the PSD. In our noise model, the noise correlation happens before the sensor converts the photons into current; thus, we consider the case of signal-dependent noise post correlation (Azzari et al., 2018)(See Eq. (53)). In our processing pipeline we first estimate, modulo a scaling factor, the noise variance function; this is done using the algorithm (Foi *et al.*, 2008). This algorithm is designed for estimating the variance function of signal-dependent white noise and when the noise is spatially correlated (i.e., not white), the algorithm’s output is off by a scaling factor that depends on the correlation (Azzari et al., 2018)(See Eq. (68)), which must be estimated separately. To identify this factor, we exploit the fact that the noise spatial correlation does not affect the statistics of an individual pixel (e.g., the variance of a pixel corresponds exactly to a point of the noise variance function). First, we estimate the mean and variance of non-overlapping 1×1×8 windows (i.e., only in the axial direction) of the 3D sequence. We discard the first and last quartiles of the measured variances to remove possible outliers. This is followed by fitting of a second affine mean-variance function over the remaining points, and comparison of the variances from the two affine models at the intensity value of the mean of the observed image: the first function is then corrected with the ratio between the two values. Next, the noise function is used to apply the generalized Anscombe transform (GAT) (Starck et al., 1998) to our data. The GAT is a variance stabilizing transformation of noisy data that makes the noise variance independent of the signal. In this way, we can estimate the noise PSD using, indiscriminately, the whole stabilized data. For practical reasons it is convenient to estimate the PSD in discrete cosine transform (DCT) domain on a small support 8×8, and then to convert it to the actual noise PSD (Azzari et al., 2018; see Eq. (77)). To estimate the PSD DCT 8×8 we first divide the whole sequence in 3D overlapping 8×8×8 cubes; then, for each cube, we first compute the 2D DCT of each 2D spatial plane, and then we apply, coefficient-wise, the squared median of absolute deviation (multiplied by 1.4826) in the axial direction: in this way we obtain a set of 2D estimates of the 2D noise PSD DCT 8×8, one for each group. Finally, we estimate the final noise 2D PSD DCT 8×8 by applying the sample median coefficientwise to all the remaining PSD estimates. Because the noise has been stabilized and a PSD estimate is now available, we can apply any of the many off-the-shelf denoising algorithms developed for correlated additive Gaussian noise to denoise our data.

We denoised the sequence using the framework introduced in (Azzari and Foi, 2016). This approach iteratively filters and refines the estimation in a multi-scale fashion, from coarse to fine scale: at each scale, it first performs binning of the data, and then it applies variance-stabilizing transformations. The resulting signal is corrupted by additive signal-independent correlated noise that can be denoised with any algorithm designed for correlated noise. This iterative approach is well suited for data affected by strong noise (such is the case of expansion microscopy), and it ensures state-of-the-art denoising results. The denoising filter that we use within the framework is the RF3D algorithm designed for videos (3D data) corrupted by correlated noise (Maggioni et al., 2014). It is important to remark that RF3D performs well in challenging cases such as 3D expansion microscopy because it exploits the redundant spatial and temporal information by promoting the sparsity of small 3D spatio-temporal volumes. The software allows the user to also perform deconvolution (deblurring) of the denoised sequence. However, the results reported here have not been deconvolved. n=2 independent experiments.

### Deconvolution

To compare the developed noise estimation, denoising, and deblurring algorithm to deconvolved data, we analyzed the image data separately after using both processing methods. The deconvolution was done with specialized deconvolution software Huygens Essential (SVI, Hilversum, Netherlands). As the ExM data is very noisy, deconvolution can be affected by artifacts if the settings are too aggressive. In our processing, SNR of 5 was used with standard strength for around 25 iterations. Compared to the pre-processed data, the deconvolved data is cleaned from noise and the features are sharp enabling the detection of the lamin network.

### SIM

Nikon N-SIM system in Nikon Ti-E inverted microscope body with CFI SR Apochromat100x/1.49 oil immersion objective and Andor iXon Ultra 897 was used for super-resolution SIM imaging. The spherical aberration correction was conducted before every imaging session by using sub-resolution fluorescent beads (PS-Speck, Thermo Fisher Scientific). This was followed by illumination pattern alignment and optimization for the used laser lines (488nm, 561nm, and 640nm). Alexa 488 and Alexa 568 were excited with 488 nm and 561 nm solid-state lasers, respectively. The SIM image reconstruction was conducted in the system by using NIS-elements software. n=2 independent experiments with 4-10 nuclei.

### ExM lamin network analysis

ExM image stacks from the 1x and 4x iterations were used to analyze the stained lamin network coverage. Overall, three areas per sample were analyzed from two samples per iteration. For this purpose, the image stacks were first divided into two sub-stacks, apical and basal. The apical sub stack was used for the final analysis. First, the stack was normalized, and a ridge detection algorithm (Steger, 1998; Wagner et al., 2017) was used to detect string-like lamin A/C structures. The following thresholded image stacks were analyzed by making a maximum intensity projection for area coverage, and by analyzing the network in Avizo 2020.2 (Thermo Fisher Scientific) for nodal and branch length analysis. First, the Auto Skeleton module was used to extract the centerlines from the network structures. Secondly, the formed graph was analyzed with the Spatial Graph Statistics module. For this analysis the n=2 independent experiments with 10 nuclei.

## Abbreviations

*ExM*: Expansion microscopy
*FWHM*: Full width at half maximum
*HSV-1*: Herpes simplex virus 1
*IT-IF*: Iterative indirect immunofluorescence staining
*LA/C-C*: Lamin A/C C-terminus
*LA/C-N*: Lamin A/C N-terminus
*LA/C-rod*: Lamin A/C rod-domain
*NE*: Nuclear envelope
*PAA*: Polyacrylamide
*SBR*: Signal-to-background-ratio
*SIM*: Structured illumination microscopy
*SR-SIM*: Super-resolution structured illumination microscopy

## Acknowledgements

This work was supported by the Academy of Finland under the award numbers 308315 and 314106 (TOI), 330896 (MVR), 332615 (EM), and the Centre of Excellence in Body-on-Chip Research (312412 (JH)), 336357 (PROFI6 - TAU Imaging Research Platform (LA, MV, AF), and by the Jane and Aatos Erkko Foundation (MVR). University of Jyväskylä Graduate School for Doctoral Studies is acknowledged for the support (SM). The authors acknowledge Biocenter Finland, and Tampere University Tampere Imaging Facility, Virus Production Facility (Title of Docent Eric Dufour), and Flow Cytometry Facility (Dr. Laura Kummola) for their services.

## Notes

### Competing Interest Statement

The authors have declared no competing interest.

### Summary of Updates

Revised Fig.2, Fig. S2, Fig. 5 Results section modified to include new results in Fig. 2 Supplemental files updated

