## Supplemental material for "Iterative immunostaining combined with expansion microscopy and image processing reveals nanoscopic network organization of nuclear lamina"

BioMediTech, Faculty of Medicine and Health Technology, Tampere University, Tampere, Finland[a]; Tampere Microscopy Center (TMC), Tampere University, Tampere, Finland[b]; Department of Biological and Environmental Science and Nanoscience Center, University of Jyväskylä, Jyväskylä, Finland[c]; Faculty of Information Technology and Communication Sciences, Computing Sciences, Tampere University, Tampere, Finland[d]; Tampere Institute for Advanced Study, Tampere University, Tampere, Finland[e]

*Equal contribution

Address: BioMediTech, Faculty of Medicine and Health Technology, Tampere, University, Tampere, Finland. Arvo Ylpön katu 34, 33520 Tampere, Finland

Characters of supplementary materials:

**Supplementary Figure 1.**

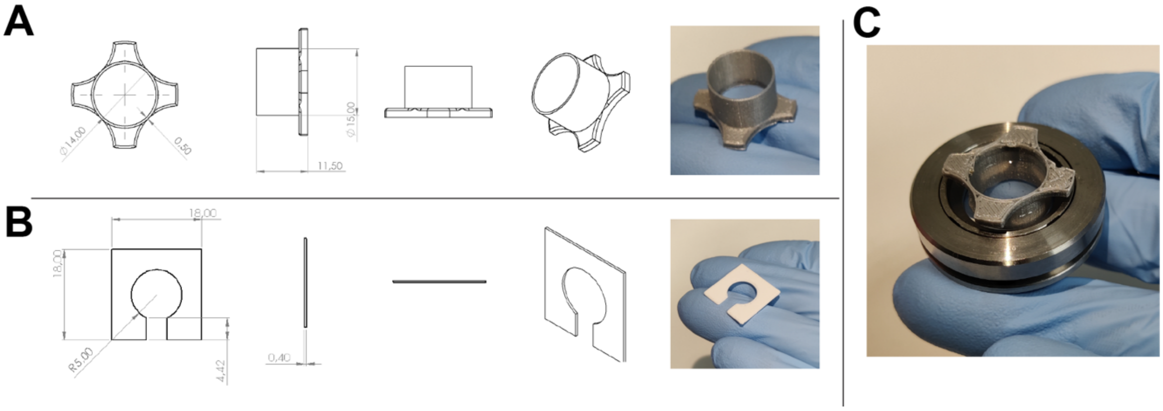


**Figure S1.** Technical tools for ExM sample preparation. In-house 3D-printed design, physical dimensions, and (A) a mold for cutting and agarose mounting of a gel piece, and (B) spacers used in applying non-polymerized gel solution to a sample specifically designed to ensure constant sample gel thickness and diameter during ExM sample preparation. (C) Gel-containing mold easily mounted with high-melt agar into an Aireka Cell for LSCM imaging.

**Supplementary Figure 2.**

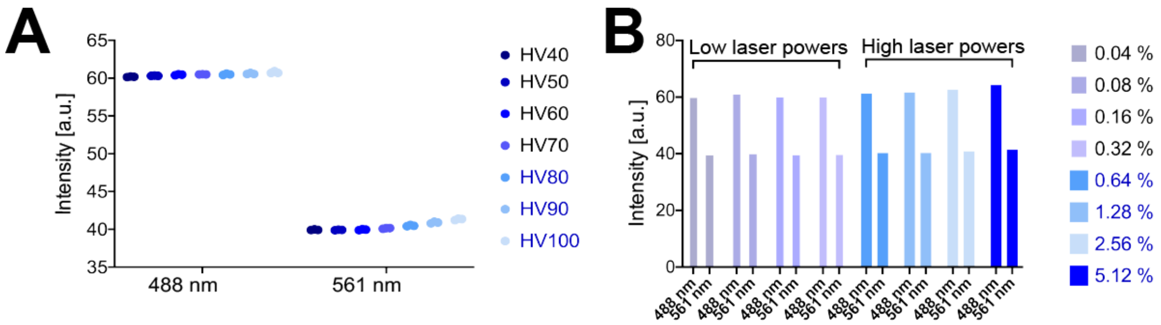


**Figure S2.** Effects of laser power and detector linearity on signal detection. (A) Detected intensities as a function of increasing detector voltages and (B) low (0.04-0.32 %) or high (0.64-5.12 %) 488 nm and 561 nm solid-state laser powers showing the linear function of the detector. n=2 individual experiments.

**Supplementary Figure 3.**

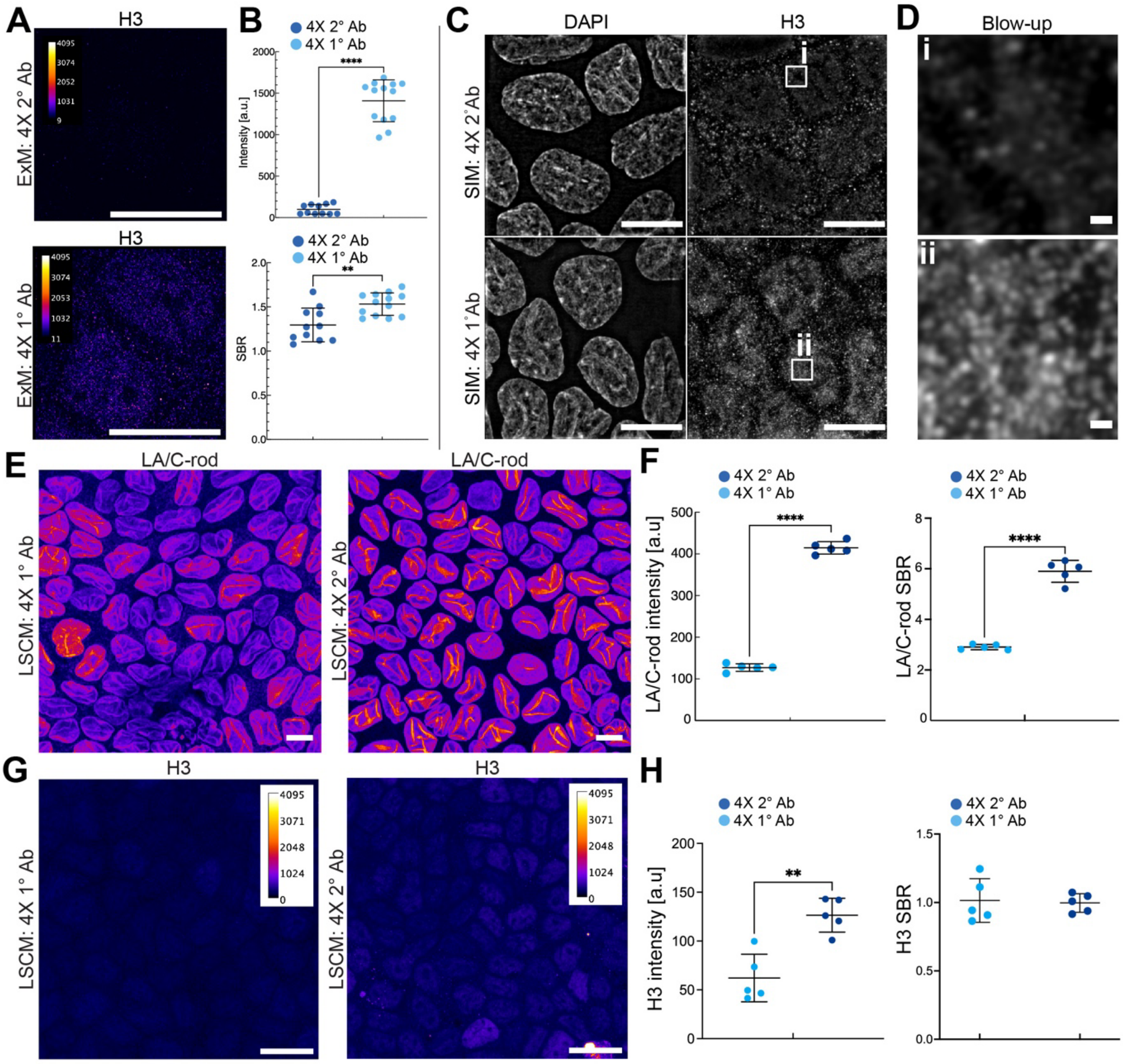


**Figure S3.** Effects of enhanced primary and secondary Ab concentration on target detection. (A) Expansion microscopy (ExM) maximum intensity Z-projection images of histone H3 using either 4X 2°Ab (upper panel) or 4X 1°Ab (lower panel) concentrations. (B) Quantifications of nuclear fluorescence intensity and signal-to-background ratio (SBR) following 4X 2°Ab and 4X 1°Ab treatments. (C) Structured illumination microscopy (SIM) maximum intensity projection images showing chromatin (DAPI) and H3 with (D) respective blow-ups (ii and ii) after treatments with 4X 2°Ab (upper panels) and 4X 1°Ab concentrations (lower panels). Scale bars, 10 μm; blow-up scale bars, 2 μm. n=2 individual experiments. (E) Representative LSCM pseudo-colored maximum intensity projection images of LA/C-rod Ab -stained epithelial nuclei using 4X 1°Ab (left panel) and 4X 2°Ab (right panel) concentrations. (F) Representative pseudo-colored intensity indicator color-coded LSCM maximum intensity projection images of H3 Ab -

stained epithelial nuclei using either 4X 1°Ab (left panel) or 4X 2°Ab (right panel) concentrations. Scale bars, 20 µm. (F) Quantification of the mean nuclear LA/C-rod fluorescence intensity and SBR in LSCM in applied Ab treatments. (G) Quantification of the mean nuclear H3 fluorescence intensity and SBR in LSCM in respective conditions. All values are presented as a mean ± standard deviation (ns $p>0.05$, *$p<0.05$, **$p<0.01$, ****$p<0.0001$, unpaired Student´s T-test).

**Supplementary Figure 4.**

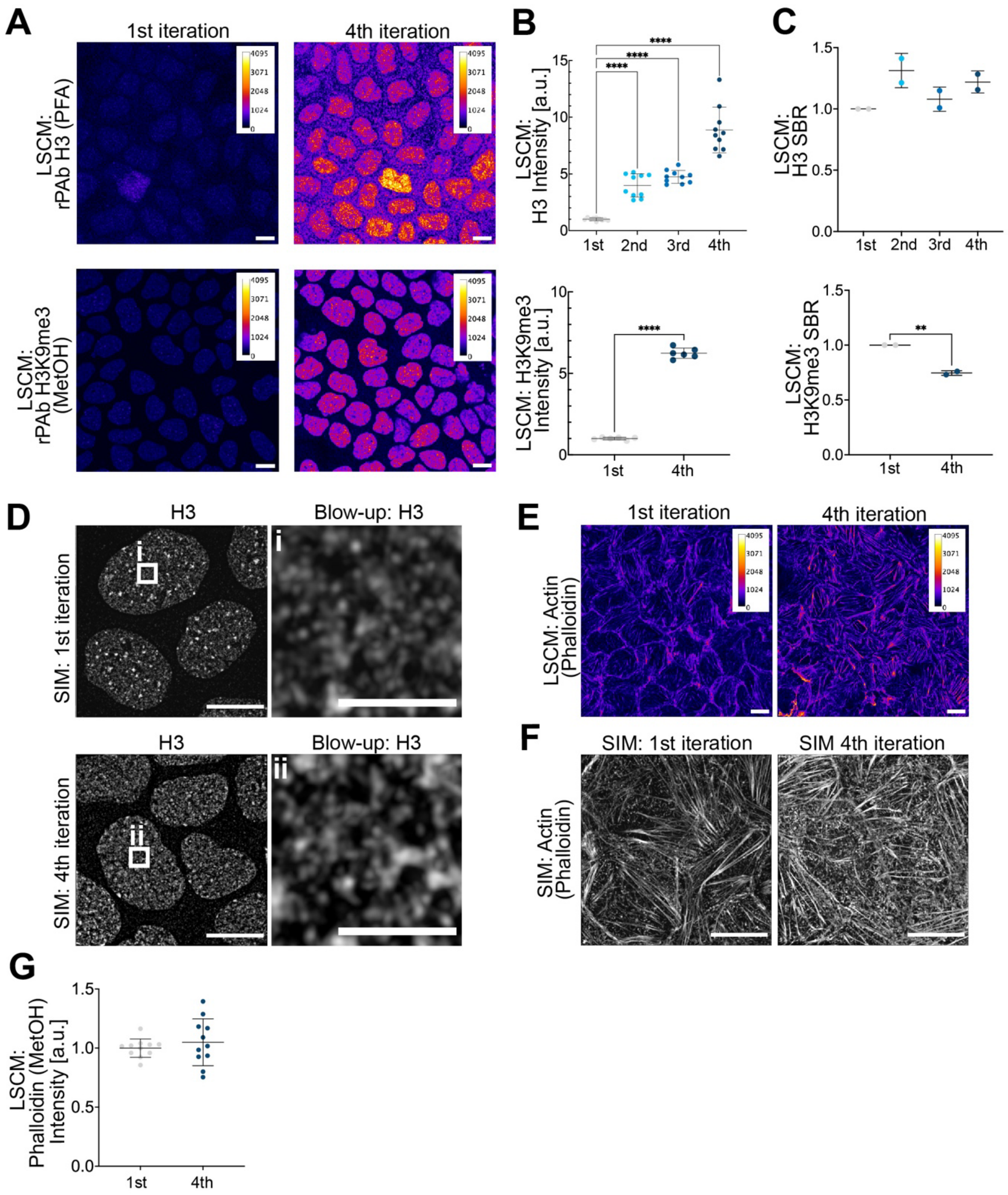


**Figure S4.** Effect of iterative immunostaining on fluorescence signal intensity and detection quality. (A) Representative LSCM pseudo-colored maximum intensity Z-projection images acquired after one (1st, far left panels) or four (4th, left panels) iterations of histone rPAb H3 (PFA fixed, upper panels) and rPAb H3K9me3 (MetOH-fixed, lower panels) staining. n=2 independent experiments. Scale bars, 10 µm. (B) Quantifications of H3 (upper panel) and H3K9me3 (lower panel) intensities in 1-4 iterations. (C)

Quantifications H3 (upper panel) and H3K9me3 (lower panel) SBRs. D) Structured illumination microscopy (SIM) images of $1^{st}$ (upper panels) or $4^{th}$ iterations (lower panels) of H3 staining and their respective blow-ups (i and ii, upper and lower right panels; scale bars, 2 µm). n=2 independent experiments. (E) Representative LSCM pseudo-colored maximum intensity Z-projection images acquired after one (left) or four (right) iterations of actin staining (MetOH fixed). (F) Representative SIM images of 1x (left) and 4x iterated (right) actin stainings (MetOH fixed). n=2 independent experiments. Scale bars, 10 µm. (G) Quantification of phalloidin intensity after 1st and 4th iterations. n=1 experiment. The data represents mean ± standard deviation. One-way ANOVA was used to test for statistical significance among multiple groups, unpaired Student´s T-test was used to test for statistical significance among two groups. ns $p>0.05$, *$p<0.05$, **$p<0.01$, ****$p<0.0001$.
